# Zfp281 inhibits the pluripotent-to-totipotent state transition in mouse embryonic stem cells

**DOI:** 10.1101/2022.03.14.484036

**Authors:** Xinpeng Wen, Zesong Lin, Hao Wu, Xudong Fu

## Abstract

The cell-fate transition between pluripotent and totipotent states determines embryonic development and the first cell-lineage segregation. However, limited by the scarcity of totipotent embryos, regulators on this transition remain largely elusive. A novel model to study the transition has been recently established, named the 2-cell-like (2C-like) model. The 2C-like cells are rare totipotent-like cells in the mouse embryonic stem cell (mESC) culture. Pluripotent mESCs can spontaneously transit into and out of the 2C-like state. We previously dissected the transcriptional roadmap of the transition. In this study, we revealed that Zfp281 is a novel regulator for the pluripotent-to-totipotent transition in mESCs. Zfp281 is a transcriptional factor involved in the cell-fate transition. Our study shows that Zfp281 represses transcripts upregulated during the 2C-like transition via Tet1 and consequentially inhibits mESCs from transiting into the 2C-like state. Interestingly, we found that the inhibitory effect of Zfp281 on the 2C-like transition leads to an impaired 2C-like-transition ability in primed-state mESCs. Altogether, our study reveals a novel mediator for the pluripotent-to-totipotent state transition in mESCs and provides insights into the dynamic transcriptional control of the transition.

## Introduction

Totipotency refers to the ability of a cell to generate all cell types[1]. In mouse embryos, zygotes and 2-cell embryos are considered totipotent cells. When the embryo develops beyond zygotes and 2-cell (2C) stages, embryos progressively lose totipotency, go through the first lineage segregation, and establish pluripotent inner cell mass (ICM) at the blastocyst stage [1]. An impairment in the totipotent-to-pluripotent state transition in mouse embryos leads to defects in embryonic development, indicating that the transition is crucial for embryonic development[2; 3; 4]. However, Limited by the material scarcity, mechanistic studies of the cell-fate transition between pluripotency and totipotency are largely impeded.

Mouse embryonic stem cells (mESCs) derived from the ICM were established as a model for pluripotency study[5]. The mESC culture can be maintained in the ground-naïve, the metastable-naïve, or the primed state[5]. When mESCs are cultured in the metastable-naïve condition, less than 1% of ESCs spontaneous transit into a totipotent-like state. This state, named as 2-cell-like (2C-like) state, exhibits several features of 2C-stage embryos, including totipotent-like developmental potential and the expression of 2C-specific transcripts, such as *Zscan4d* and MERVL repeats [6; 7].

The 2C-like transition is initiated by the transcription factor Dux. After Dux activation, pluripotent mESCs transits into the 2C-like state (the entry of 2C-like transition). The 2C-like state is unstable, and 2C-like cells can spontaneously transit back to the pluripotent state (the exit of the 2C-like transition). Notably, the 2C-like transition recapitulates the transcriptomic features of the transition between totipotency and pluripotency of embryonic development[8; 9]. Thus, the 2C-like transition is currently a widely-used model for mechanistic exploration of cell-fate transition between totipotency and pluripotency[2; 3; 4; 7; 8; 9; 10; 11; 12; 13; 14; 15; 16; 17; 18; 19; 20; 21; 22; 23; 24; 25; 26; 27; 28; 29; 30].

Our previous studies generated an inducible 2C-like transition model, a reporter mESC cell line containing a MERVL-promoter-driven reporter, and a doxycycline-inducible Dux transgene (synDux)[7]. The synDux can drive the pluripotent-to-2C-like transition in mESCs, and the reporter can indicate whether the cells are in the 2C-like state. Importantly, synDux-induced 2C-like transition recapitulates the spontaneous 2C-like transition in the mESC culture[7; 8]. Thus, this cell line is a valuable tool to monitor the entry and exit of the 2C-like transition. Using this model, we constructed the comprehensive roadmap for the 2C-like transition and revealed the regulatory network controlling the transition via a genome-wide CRISPR-Cas9-mediated screen[7; 8].

By examining the screen result, we identified that Zfp281 is a candidate factor affecting 2C-like transition. Zfp281 is a transcription factor modulating cell-fate transitions through transcriptional regulation. For instance, Zfp281 inhibits the expression of naïve-pluripotent-related genes via the interaction of Tet1 and consequentially promotes the naïve-to-prime transition in mESCs[31]. In addition, Zfp281 inhibits the transition from pluripotent cells to extraembryonic endoderm stem cells (XENs) by interacting with polycomb repressive complex 2 (PRC2)[32]. Furthermore, Zfp281 promotes mESCs to transit into trophoblast stem cells (TSCs) via recruiting complex proteins associated with Set1 (COMPASS) to activate TSC-related genes[33]. Taken together, these results suggest that Zfp281 plays a critical role in the cell-fate transition in mESCs. To this end, we set out to examine the function of Zfp281 in the 2C-like transition.

## Materials and Methods

### ESC culture

The mESC-E14 cell line with MERV-L-LTR-tdTomato reporter was kindly provided by the laboratory of Jin Zhang from Zhejiang University. To generate the inducible 2C-like transition model, this reporter cell line was infected with lentivirus containing inducible dux (addgene, cat. no. 138320), and clones were picked. The mESCs with the 2C::tdTomato reporter and doxycycline-inducible codon-optimized Dux cells were cultured on 0.1% gelatin-coated plates with standard LIF/serum medium containing 10% FBS (HAKATA, cat. no. HB-FBS-500), 1,000 U ml–1 mouse LIF (Biolegend, cat. no. 554002), 0.1 mM non-essential amino acids (Gibco, cat. no. 11140), 0.055 mM β-mercaptoethanol (Gibco, cat. no. 21985023), 2 mM GlutaMAX (Gibco, cat. no. 35050), 1 mM sodium pyruvate (Gibco, cat. no. 11360) and penicillin/streptomycin (100 U ml–1) (Gibco, cat. no. 15140) in a humidified 5% CO2 atmosphere at 37°C. For the culture of ESC lines, the medium was changed daily, and cells were routinely passaged every other day.

### FACS

Flow cytometry analysis was performed using the BD FACSAria Fusion SORP. Data and images were analyzed and generated using FlowJo (V10) software. The gating strategy was shown in FACS figures.

### RNA isolation and qPCR

Cellular RNA was collected using the FastPure Cell/Tissue Total RNA Isolation Kit V2 (Vazyme, cat. no. RC112). Complementary DNA was generated using the HiScript II 1st Strand cDNA Synthesis Kit (+gDNA wiper) (Vazyme, cat. no. R212), and qRT-PCR was performed using the Taq Pro Universal SYBR qPCR Master Mix x (Vazyme, cat. no. Q712) on XN-1000V (ABI). Relative quantification was performed using the comparative CT method with normalization to *Gadph*. Primers used in the qPCR are listed in Table S1.

### Construction of siRNA and cell transfection

Two pairs of interference sequences that targeted mouse Tet1, Cxxc1, Kmt2d mRNA were designed and synthesized using the siRNA online design program (Merck). The three interference sequences were Tet1-1(5’-tgtagaccatcactgttcgac-3’), Tet1-2(5’-gagattaacgctggaacaag-3’), Cxxc1-1(5’-gtcgcaaaaccggacatcaatt-3’), Cxxc1-2(5’-acgagcttgaggccatcattc-3’), and Kmt2d-1(5’-gttcatcgagttgcgacataa-3’), Kmt2d-2(5’-gtcctataaccagcggagtct-3’). DNA oligos containing the target sequences were annealed and synthesized by T7 RNAi Transcription Kit (Vazyme, cat. no. TR102-01). Transfection was mediated by Lipo6000 transfection reagent (Beyotime, cat. no. C0526). The cells post transfection was then collected for other assays.

### CRISPR–Cas9

The gene knockdown by CRISPR–Cas9 was performed in our previous reports[7]. The sgRNA sequences are listed in Table S2. Lentivirus was produced using the psPAX2-PMD2.G system in 293T cells. The mESCs were infected with lentivirus for 48 h in a medium containing1 μg/ ml Polybrene. After two days of infection, cells were cultured in a medium containing 1 μg /ml puromycin for another eight days to select for infected cells.

### Western blotting

Cellular protein was purified using RIPA Lysis Buffer with protease inhibitor. Western blotting was carried out with gradient gel (BCM Biotech, cat. no. P2012) with the following antibodies: Zfp281 (1:3,000, ABclonal, cat. no. A12650), GAPDH (1:20,000, ABclonal, cat. no. AC033), goat anti-rabbit IgG (H+L) secondary antibody, HRP (1:10,000, ABclonal, cat. no. AS014) and goat anti-mouse IgG (H+L) secondary antibody, HRP (1:10,000, ABclonal, cat. no. S003).

### Pseudo-genome preparation

As repeat elements tend to have multiple highly similar copies along the genome, it is relatively complex to accurately align them and estimate their expression. Hence, we created a repeat pseudo-genome. We used a slightly modified version of the RepEnrich (v0.1)[34] software. Briefly, for each repetitive element subfamily, a pseudo-chromosome was created by concatenating all genomic instances of that subfamily along with their flanking genomics 15bp sequences and a 200bp spacer sequence (a sequence of Ns). The pseudo-genome was then indexed using STAR (v.2.5.2b)[35], and the corresponding gtf and refFlat files were created using custom scripts and by considering each pseudo-chromosome as one gene.

### Sequencing alignment for coding genes

Raw reads were first trimmed using Trimmomatic (v.0.36). Illumina sequence adaptors were removed, the leading and tailing low-quality base pairs (fewer than 3) were trimmed, and a 4-bp sliding window was used to scan the reads and trim when the window mean quality dropped below 15. Only reads having at least 50-bp were kept. The resulting reads were mapped to the mm10 genome using STAR[35] (v.2.5.2b) with the following parameters: -outSAMtype BAM SortedByCoordinate –outSAMunmapped Within –outFilterType BySJout -outSAMattributes NH HI AS NM MD -outFilterMultimapNmax 20 -outFilterMismatchNmax 999 -quantMode TranscriptomeSAM GeneCounts. The generated gene expression count files generated by STAR were then used for estimating gene expression.

### Sequencing alignment for repeats

Multi-mapped reads and reads mapping to intronic or intergenic regions were extracted and then mapped to the repeat pseudo-genome. First, the TagReadWithGeneExon command of the dropseq tools (v1.13)[36] was used to tag the reads into utr, coding, intergenic and intronic reads using the bam tag ‘XF’. Multi-mapped reads, intergenic and intronic reads were extracted and mapped to the repeat pseudo-genome using STAR. The STAR read counts were used as an estimate of repeat expression.

### RNA-seq normalization

For each sample, the gene and repeat expression matrices were merged. Then the ‘Trimmed Mean of *M* values’ normalization (TMM) method[37] from the R/Bioconductor package edgeR package (v3.24.0) was used to calculate the normalized expression[38].

### Differential gene expression analysis of bulk RNA-seq data

The R/Bioconductor edgeR package (v3.24.0)[38] was used to detect the differentially expressed genes between the different samples using the generalized linear model-based method. Genes showing more than twofold expression change and an FDR<0.0001 were considered as differentially expressed.

### Functional enrichment analysis

Clusterprofiler was used to perform GO function enrichment and KEGG pathway annotation[39]. The associated GO and pathway enrichment plots were generated using the ggplot2 package (v3.1.0).

### ChIP-seq data analysis

The Chip-seq data were downloaded from GEO datasets, and below are the corresponding GEO Accession numbers: GSE81045 (Zfp281), GSE24843 (Tet1), GSE12721 (H2K119ub), GSE158460 (H3K9me3), GSE48519 (H3K4me1, H3K4me3). Raw reads were trimmed using Trimmomatic53 and then mapped to the mm10 genome using Bowtie254 (v2.2.9). Multi-mapped and unmapped, low-quality reads were removed using sambamba55 (0.6.6). Chip-seq peaks were determined by MACS (v2.0.10), and the peaks were visualized using IGV software.

## Results

### The transcription factor Zfp281 inhibits the pluripotent-to-2C-like transition

Our previous screen results indicate that Zfp281 regulates the 2C-like transition[7] (Fig. 1A). To validate the hypothesis, we used the inducible 2C-like transition model (Fig. S1A) to examine the role of Zfp281 on the 2C-like state. We design two independent sgRNA targeting Zfp281 and verify their efficiency (Fig. 1B). Our results show that Zfp281-perturbation significantly increases the population of 2C-like cells and the expression of 2C-like-state marker genes after 24-hour synDux induction (Fig. 1C–D). Importantly, Zfp281 perturbation also increases the spontaneous 2C-like transition, and the expression of synDux is not altered upon Zfp281 perturbation (Fig. S1B-C), indicating that Zfp281 does not mediate 2C-like transition through synDux. Altogether, these results suggest that Zfp281 regulates the 2C-like transition.

**Fig. 1.**
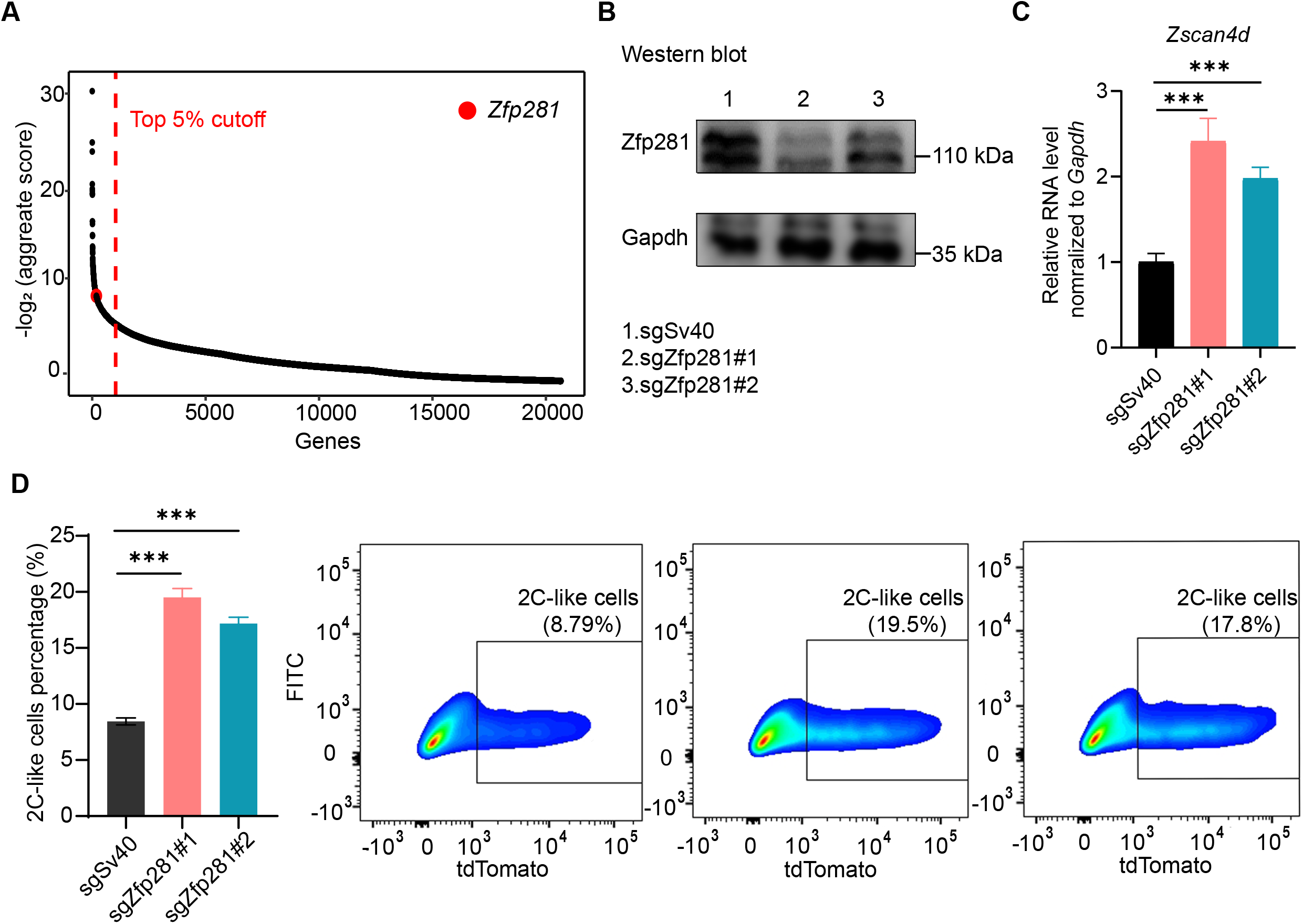
Zfp281 regulates the 2C-like transition of mESCs. (A) The sgRNA count enrichment from the CRISPR screen. Zfp281 is one of the top candidates mediating the 2C-like transition in the screen. (B) Western-blot confirming efficient Zfp281 perturbation. (C) Relative *Zscan4d* mRNA levels normalized to *Gapdh* in *Dux*-activated mESCs. (D) The percentage of 2C-like cells after 24h *Dux* induction of the indicated manipulation in mESCs by FACS. (B-D) The X in sgX refers to the gene that sgRNA targets to; sgSv40 is negative control. Shown are mean ± s.d, P values were calculated by unpaired t-test, two-tailed, two-sample unequal variance, *** < 0.001.

Zfp281 may affect the initiation, the entry, or the exit of the 2C-like transition. We find that Zfp281-perturbation does not affect Dux expression (Fig. S1D), and Zfp281-perturbed cells without synDux induction exhibit minimal changes on the transcripts upregulated or downregulated during the 2C-like transition (these transcripts are named as 2C-upregulated/downregulated transcripts respectively in the following text, Fig. S1E and Table S3). These results suggest that Zfp281 does not affect the initiation of 2C-like transition. In addition, our results show that Zfp281-perturbed cells exhibit similar maintenance of the 2C-like state (Fig. S1F), implying that Zfp281 does not modulate the exit of 2C-like transition. Taken together, these results suggest that Zfp281 impedes the entry of 2C-like transition.

### Zfp281 inhibits the expression of 2C-upregulated transcripts during the 2C-like transition

Previous studies indicated that Zfp281 regulated cell-fate transition through transcriptional regulation[31; 32; 33; 40; 41; 42]. Thus, to understand how Zfp281 affects the 2C-like transition, we focus on the transcriptional effect of Zfp281 on the 2C-like transition. We perform RNA-seq on Zfp281-perturbed cells and control cells after 24-hour synDux induction (Fig.S2A-B). By comparing the transcriptome, we identify 1081 upregulated, and 407 downregulated transcripts upon Zfp281 perturbation (Fig. 2A and Table S3). The upregulated genes/repeats are Zfp281-repressed transcripts during the 2C-like transition. They include 2-cell-embryo-specific transcripts[6] such as *Zscan4d, Zfp352*, and MERVL-int (Fig. 2B), and the GO term of these transcripts is enriched in cell fate commitment (Fig. S2C). Importantly, most of these Zfp81-repressed transcripts are 2C-upregulated transcripts (Fig. 2C), further supporting that Zfp281 impedes the pluripotent-to-2C-like transition. The downregulated genes include *Dnmt3a, Eif2s3y*, *Lefty2* (Fig. 2B) and are functionally enriched in Glycine, serine, and threonine metabolism (Fig. S2C).

**Fig. 2.**
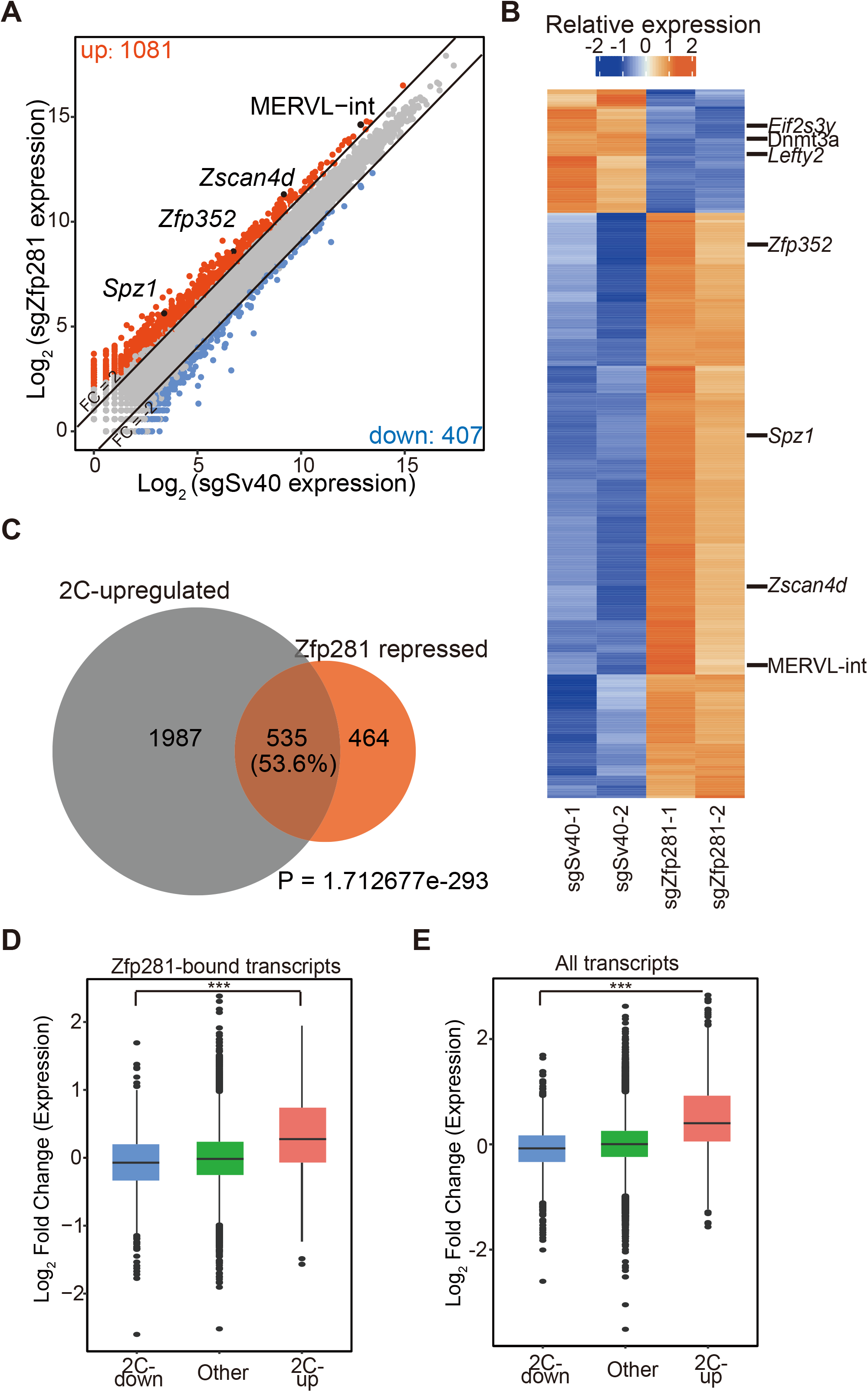
Zfp281 impedes the expression of 2C-upregulated transcripts during the 2C-like transition. (A) A scatter plot comparing the gene/repeat expression profiles between control and Zfp281-perturbed mESCs after *Dux* induction. The criteria for gene changes are Fold change (FC) > 2 and False discovery rate (FDR) < 0.05. (B) A heatmap showing the relative expression levels of differential expression genes/repeats (FC > 1, FDR < 0.05) from two biologically independent samples of control and Zfp281-perturbed mESCs after *Dux* induction. (C) A Venn diagram demonstrating overlaps of Zfp281-repressed transcripts and 2C-upregulated transcripts. Zfp281-repressed transcripts are defined as the transcripts activated in Zfp281-perturbed mESCs compared to control mESCs after *Dux* induction (FC > 2 and FDR < 0.05). P value was calculated by a hypergeometric test. (D) A box plot showing the log_2_ (FC) of Zfp281-bound transcripts after *Dux* induction. Within all Zfp281-bound transcripts, 2C-upregulated transcripts exhibit the most significant changes. (E) A box plot showing the expression levels of 2C-regulated and other transcripts after *Dux* induction. (A, B) FDRs were estimated using the Benjamini–Hochberg method on the P values of the two-sided quasi-likelihood F-test calculated using the edgeR package. (D, E) The black central line is the median value, the box limits indicate the upper and lower quartiles. (A-E) The X in sgX refers to the gene that sgRNA targets to; sgSv40 is negative control. (D-E) P values were calculated with the absolute value of the log2 (FC) by the Wilcoxon rank-sum, *** < 0.001. The dots represent outliers.

To identify how Zfp281 shapes the transcriptome of 2C-like transition, we compare the ChIP-seq results of Zfp281 with the RNA-seq data[31]. We find that although Zfp281 can bind to both 2C-downregulated and 2C-upregulated genes (Fig. S2D), the transcriptional changes of Zfp281-bound-2C-upregulated transcripts are more significant than that of Zfp281-bound-2C-downregulated transcripts (Fig. 2D). Additionally, the change of total 2C-upregulated genes is significantly higher than that of total 2C-downregulated genes upon Zfp281 perturbation (Fig. 2E), suggesting that Zfp281 mainly affects 2C-upregulated genes during the 2C-like transition. Interestingly, we find that Zfp281-unbound 2C-upregulated genes are significantly increased upon Zfp281 perturbation, further implying that the transcriptional effect of Zfp281 promotes the 2C-like transition (Fig. S2E).

### Tet1 mediates the inhibitory effect of Zfp281 on the 2C-upregulated genes

Zfp281 shapes transcriptome through recruiting epigenetic elements to targeted genes[31; 32; 33; 40; 41; 42]. For instance, Zfp281 inhibits the transcription of naïve-pluripotent genes by binding with Tet1 in mESCs[31]. To search for the factors that contributed to the effect of Zfp281 on 2C-upregulated transcripts, we analyze the ChIP-seq data of epigenetic factors reported to interact with Zfp281[33; 42]. We find that Zfp281 colocalizes with H3k4me1, H3k4me3, and Tet1 in mESCs (Fig. S3A) but shows no colocalization of H2AK119ub and H3K9me3. Based on the ChIP-seq data, we chose two factors mediating H3K4 methylation (Cxxc1 and Kmt2d) and Tet1 for further study.

Firstly, we focus on the methylation of H3K4. We design two siRNA targeting *Cxxc1* and find that these two siRNA do not consistently affect the expression of *Zscan4d* and MERVL upon Dux activation (Fig. S3B). Additionally, two siRNA targeting *Kmt2d* show similar effects on *Zscan4d* and MERVL (Fig. S3B). These results indicate that the methylation of H3K4 does not affect 2C-upregulated genes during the 2C-like transition.

We next focus on Tet1. The major role of Tet1 is DNA demethylation[43]. Interestingly, Tet1 plays a dual role in shaping the transcriptome of mESCs[43]. It can activate and inhibit gene transcription by interacting with distinct epigenetic factors.Tet1 directly interacts with Zfp281 in mESCs[31]. In addition, the Tet family participates in the regulation of the 2C-like transition [23; 24; 32]. All these suggest that Tet1 contributes to the inhibitory effect of Zfp281 on 2C-upregulated transcripts during the 2C-like transition. To validate the hypothesis, we analyze the Tet1 and Zfp281 ChIP-seq data in mESCs. The majority of Zfp281-bound 2C-upregulated genes are bound by Tet1(Fig. 3A). Furthermore, within Zfp281-bound 2C-upregulated genes, the ones that are bound by Tet1 exhibits higher transcriptional changes than those not bound by Tet1 upon Zfp281 perturbation (Fig. S3C-E), suggesting that Tet1 mediates the inhibitory effect of Zfp281 on the 2C-upregulated genes.

**Fig. 3.**
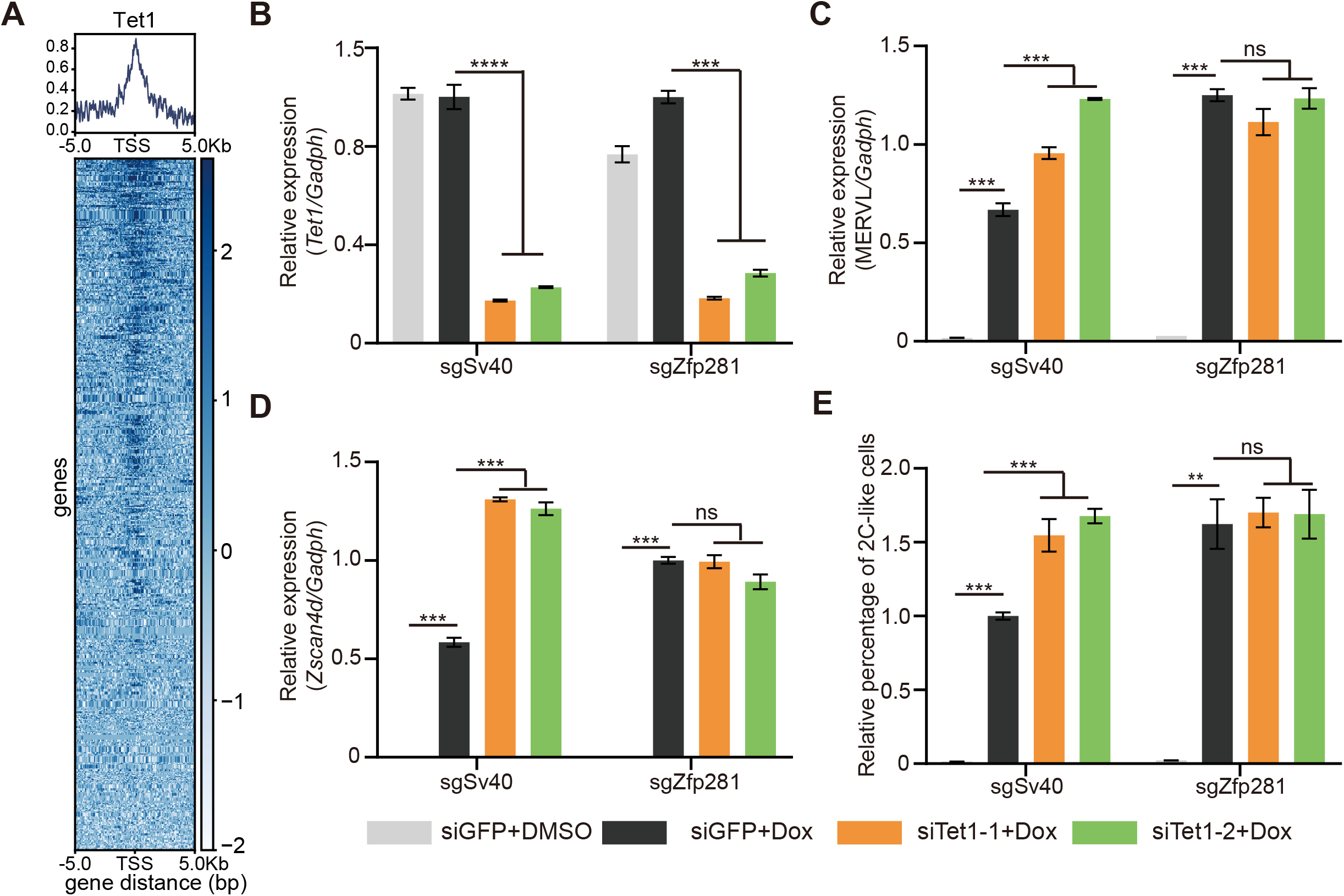
Zfp281 represses the 2C-like transition through Tet1. (A) Average occupancy plots and heatmaps of Tet1 signal within 5 kb of the center of TSS (transcription start sites) region of the Zfp281-bound-2C-upregulated genes. (B-D) Relative mRNA levels of *Tet1* (B), MERVL (C), and *Zscan4d* (D) normalized to *Gadph* in mESCs upon indicated manipulation. (E) The relative percentage of 2C-like cells in mESCs upon indicated manipulation. (B-E) The siTet1-1/2 represents two independent siRNA targeting Tet1, siGFP is the negative control. Dox represents doxycycline. The X in sgX refers to the gene that sgRNA targets to; sgSv40 is negative control. Shown are mean ± s.d, n = 3. P values were calculated by unpaired t-test, two-tailed, two-sample unequal variance, ns = no significance, ** < 0.01, *** < 0.001, **** < 0.0001.

To further confirm that Tet1 mediates the effect of Zfp281 on 2C-upregulated genes, we knockdown Tet1 in Zfp281-perturbed and control mESCs. After *Dux* induction, Tet1 deficiency significantly increased *Zscan4d* and MERVL expression while showing no effect on these genes in Zfp281-perturbed cells (Fig. 3B–D). Our results suggest that Tet1 mediates the suppression effect of Zfp281 on 2C-upregulated genes during the 2C-like transition.

Lastly, we find that Tet1 mediates the inhibitory effect of Zfp281 on the 2C-like transition (Fig. 3E). Tet1 knockdown significantly facilitates the 2C-like transition in control cells but does not affect the 2C-like transition in Zfp281-perturbed cells. This result not only suggests that Zfp281 inhibits the 2C-like transition via Tet1 but also supports that Zfp281 mediates the 2C-like transition through transcriptional regulation.

### Zfp281 contributes to the impaired 2C-like-transition ability in the primed-state mESCs

There are three major well-studied pluripotent states, which are ground-naïve state, metastable-naïve state, and primed state[5]. The mESCs in the ground-naïve state exhibit significantly lower 2C-like transition compared to mESCs in the metastable-naive state[6]. One of the reasons is that ground-naïve mESC exhibits a higher expression of Nanog, which inhibits the 2C-like transition[15]. The 2C-like transition in primed pluripotency has not been investigated. Interestingly, primed-state mESC shows higher Zfp281 expression compared to that of naïve-state mESCs[31]. Given that Zfp281 inhibits the 2C-like transition, we hypothesized that primed-state mESCs might exhibit decreased potential for the 2C-like transition.

To validate the hypothesis, we firstly compare the 2C-like transition ability of mESCs cultured in ground-naïve, metastable-naïve, and primed states. Consistent with previous results, ground-naïve state mESCs exhibit impaired 2C-like transition ability than mESCs cultured in the metastable-naïve state. Interestingly, primed-state mESCs exhibit an even lower ability of 2C-like transition compared to that of ground-naïve state mESCs (Fig. 4A).

**Fig. 4.**
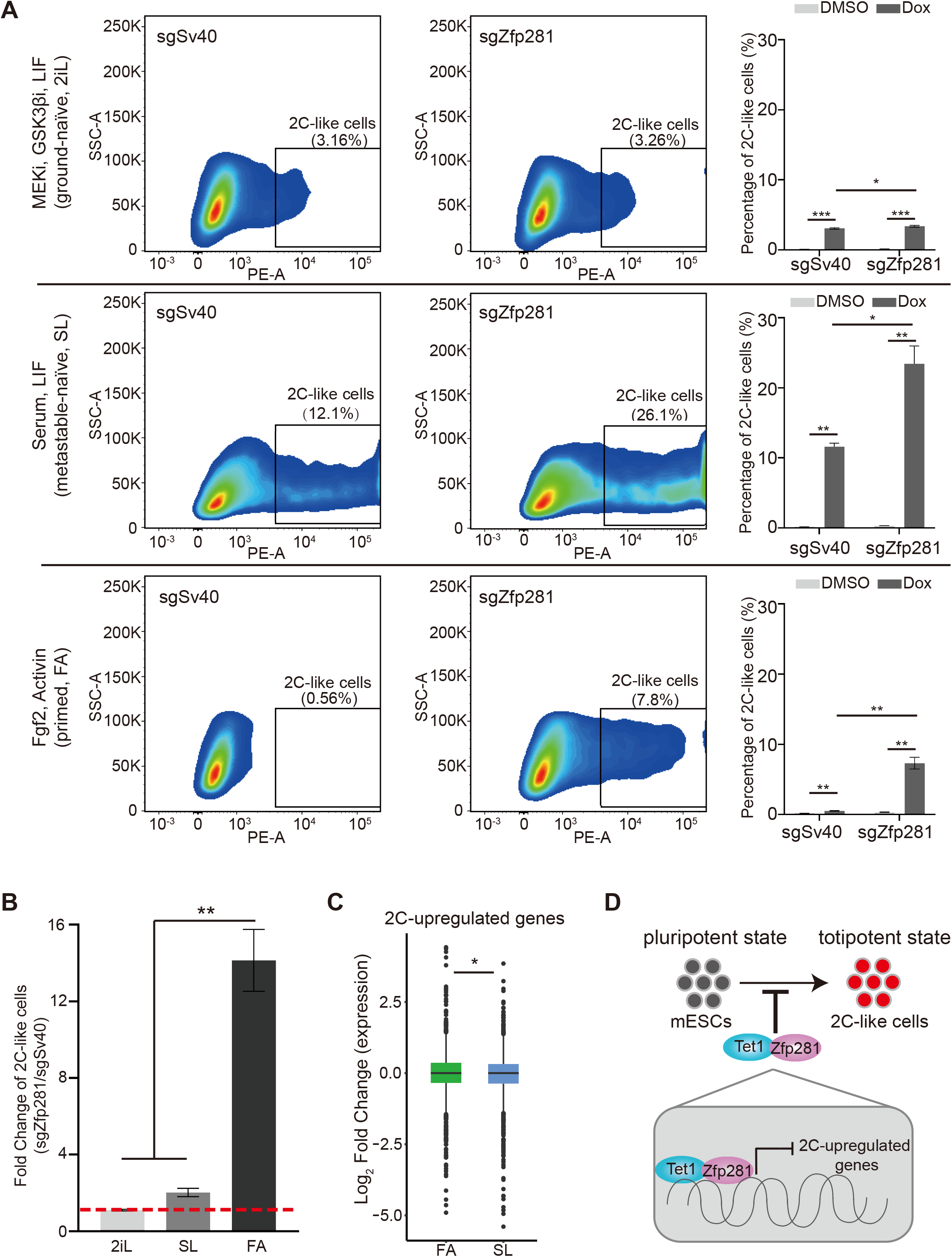
Zfp281 leads to an impaired 2C-like-transition ability in primed-state mESCs. (A) The percentage of 2C-like cell population in ground-naïve (2iL), metastable-naïve (SL), and primed states (FA) mESCs upon indicated perturbation. Representative FACS results are shown. (B) The fold change of the 2C-like cell population in three pluripotent states Zfp281-perturbed mESCs normalized to control mESCs. Redline indicates fold change of 1. (A-B) Shown are mean ± s.d, n = 3. P values were calculated by unpaired t-test, two-tailed, two-sample unequal variance, * < 0.05, ** < 0.01, *** < 0.0001. (C) A box plot demonstrating the log_2_ (fold change) of 2C-regulated genes expression in metastable-naïve and primed states mESCs upon Zfp281 perturbation. P values were calculated by the Wilcoxon rank-sum test, * < 0.05. The dots represent outliers. (D) A model showing Zfp281 and Tet1 contributes to the impaired 2C-like-transition ability by repressing the expression of 2C-upregulated genes. (A-C) Dox represents doxycycline. The X in sgX refers to the gene that sgRNA targets to; sgSv40 is negative control.

To test whether Zfp281 contributes to the decreased 2C-like transition in primed-state mESCs, we compare the effect of Zfp281-perturbation on 2C-like transition in mESCs maintained in different states. Although Zfp281 deficiency increases the 2C-like cells population in each pluripotent state (Fig. 4A), the effects are distinct. The impact of Zfp281 on the 2C-like transition is most significant in primed-state mESCs (Fig. 4B). The transcriptomic analysis also suggests that Zfp281-perturbation has a more substantial effect on 2C-upregulated transcripts in primed-state mESCs than in naïve-state mESCs (Fig. 4C). On the contrary, Zfp281 exhibits a marginal impact on the 2C-like transition in ground-naïve-state mESCs (Fig. 4A–B), which is consistent with the low expression of Zfp281 in ground-naïve state mESCs. These results indicate that Zfp281 contributes to the decreased 2C-like transition in primed-state mESCs but does not play a major role in the reduced 2C-like transition in ground-naïve-state mESCs.

## Discussion

The cell-fate transition between pluripotent and totipotent states is crucial for embryonic development. However, it is challenging to examine the transition due to the limited relevant biological materials. The 2C-like transition in mESCs has recently become a novel model to study the transition[9; 30]. In this study, by using an inducible 2C-like transition model, we revealed that Zfp281 impedes the pluripotent-to-2C-like transition (Fig. 4D). Mechanistic-wise, we showed that Zfp281 inhibits the activation of 2C-upregulated genes through the interaction with Tet1.

Previous results indicate that Zfp281 plays an important role in mediating cell-fate-transition, including the transition between naïve and primed state pluripotency. Here, we revealed a novel function of Zfp281 in mediating pluripotent-to-2C-like state transition, further supporting that Zfp281 is a master regulator for cell-identity determination.

The 2C-like transition is initiated by the transcription factor Dux and is reversible. Our results indicate that Zfp281 specifically impedes the entry of 2C-like transition but exhibits no effect on the initiation or the exit of 2C-like transition. Notably, we found that Zfp281 shows no impact on the maintenance of the 2C-like state, indicating that Zfp281 is not required for the self-renewal of the 2C-like state.

It has been reported that the Tet family inhibits the 2C-like transition by maintaining the expression of pluripotent genes, but the individual effect of the Tet family member on the 2C-like transition has not been carefully examined[24]. In this study, we showed that Tet1 interacts with Zfp281 and inhibits the 2C-like transition through impeding the activation of 2C-upregulated genes, indicating that the Tet family plays multi-dimensional roles in the pluripotent-to-totipotent transition.

Lastly, our study compared the potential for the 2C-like transition in ground-naïve, metastable-naïve, and primed state pluripotent stem cells. Our results revealed that primed-state mESCs exhibit decreased potential for 2C-like transition, and Zfp281 contributes to the decrease. On the contrary, Zfp281 plays a minimal role in the 2C-like transition in ground-naïve-state mESCs, suggesting the role of Zfp281 on the 2C-like transition is dependent on the pluripotent state.

In conclusion, our study reveals the function of Zfp281 on the 2C-like transition and the underlying mechanisms. It is interesting to investigate whether Zfp281 plays a similar role in totipotent mouse embryos and human mESCs.

## Author Contributions

Xudong Fu contributed to the study conception and design. Material preparation, data collection, and analysis were performed by Xudong Fu, Xinpeng Wen, Zesong Lin, and Wu Hao. The first draft of the manuscript was written by Xudong Fu, Xinpeng Wen, and Zesong Lin. All authors commented on previous versions of the manuscript, read, and approved the final manuscript.

## Acknowledgments

We thank the technical support by the Core Facilities, Liangzhu Laboratory; Dr. Jin Zhang, and Dr. Li Shen for their help in establishing the reporter cell line. Xudong Fu is supported by the National Key Research and Development Program of China (2021YFC2700101) and the hundred talents program of Zhejiang University.

## Data Availability Statement

All data generated or analyzed during this study are included in this published article and its supplementary information files. The RNA-seq dataset generated or analyzed during the current study has been deposited to NCBI Gene Expression Omnibus (GEO). All codes used during this study are available upon reasonable request.

