## supplement figure for "Zfp281 inhibits the pluripotent-to-totipotent state transition in mouse embryonic stem cells"

### *Supplementary Material*

#### Supplementary Figures

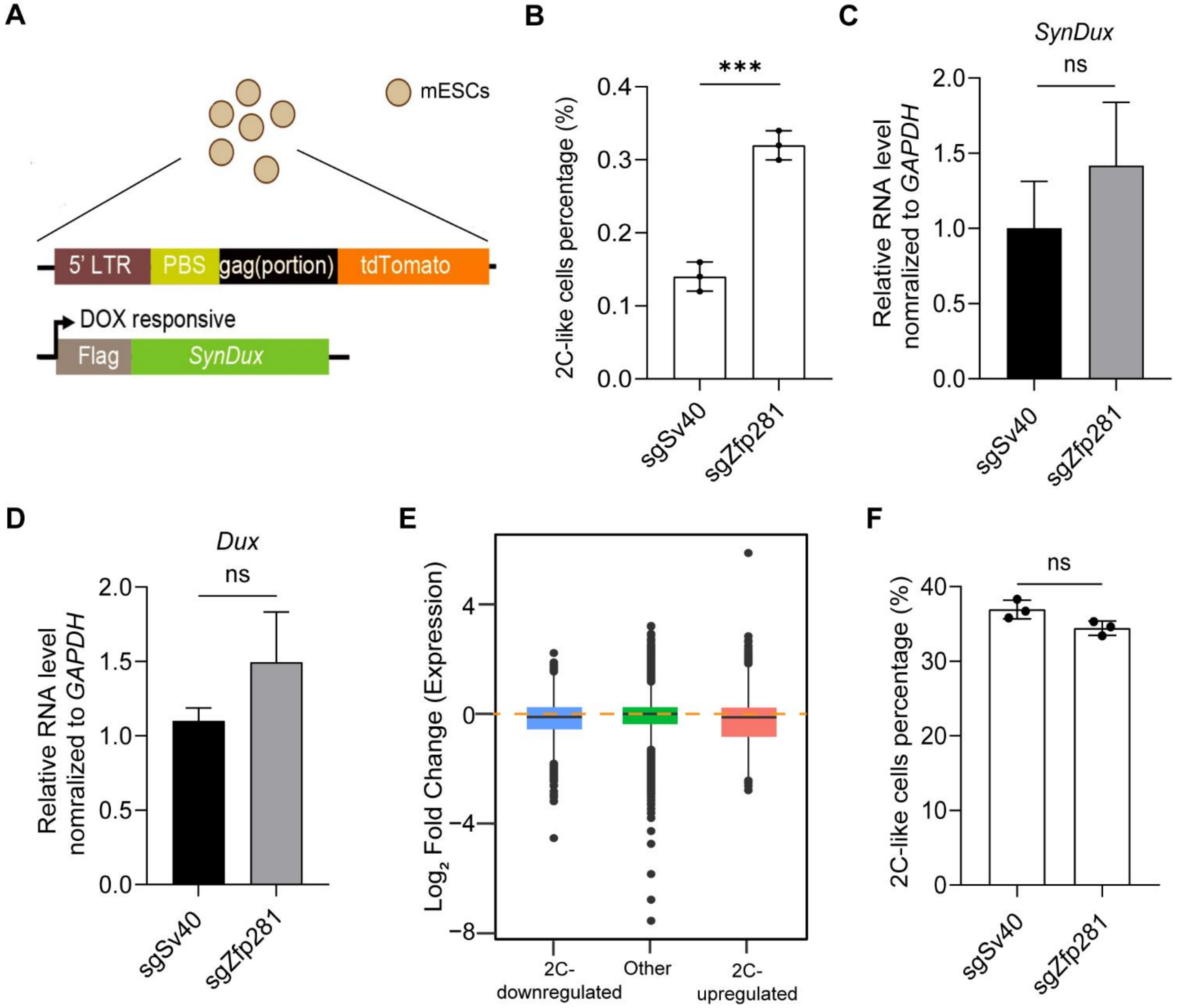


**Fig. S1, Zfp281 inhibits the pluripotent-to-2C-like state transition in mESCs.** (A) A schematic representation of the constructs in the mESCs. tdTomato is under the control of the MERVL promoter. syn*Dux* refers to codon-optimized exogenous *Dux*. PBS, primer binding site. LTR, long terminal repeats. (B) The percentage of spontaneous 2C-like cells of the indicated manipulation in three independent mESCs by FACS. (C, D) Relative *synDux* and *Dux* mRNA levels normalized to *Gapdh* in Dux-activated mESCs. (E) A box plot showing the $\log_2$ (fold change) of 2C-regulated and other genes in Zfp281-perturbed normalized to control mESCs before *Dux* induction. The black central line is the median, the box limits indicate the upper and lower quartiles. The dots represent outliers. The orange dashed line indicates a fold change of 1. (F) Flow cytometry results of the maintenance of the 2C-like state. (B, C, D, F) The X in sgX refers to the gene that sgRNA targets to; sgSv40 is negative control. Shown are mean ± s.d, n = 3. P values were calculated by unpaired t-test, two-tailed, two-sample unequal variance, ns = no significance, *** < 0.001.

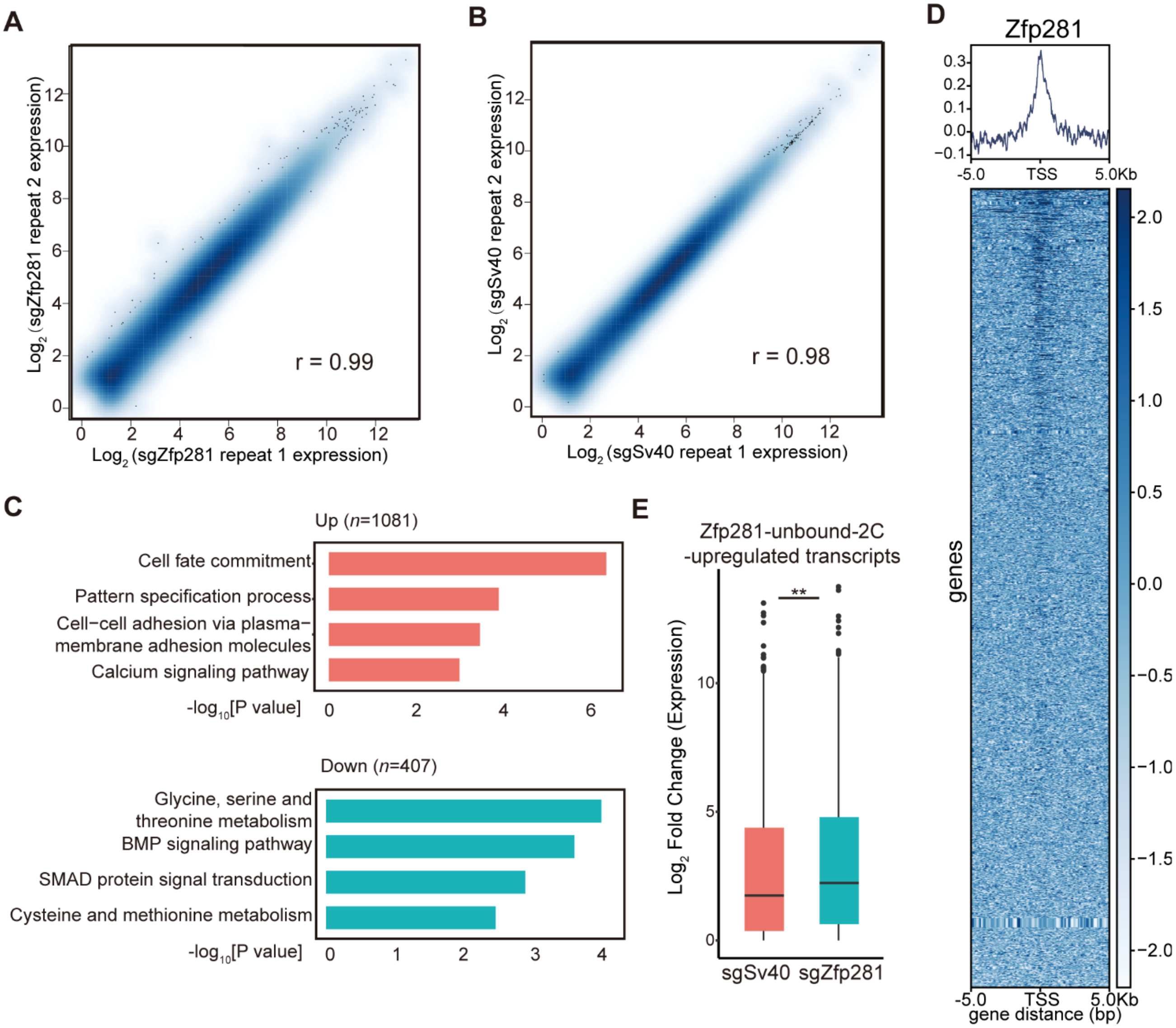


**Fig. S2, Zfp281 regulates the transcriptome of the 2C-like transition.** (A) Scatter plot comparing the gene expression profiles between two biologically independent samples of *Dux*-activated Zfp281-perturbed mESCs (Pearson correlation, r = 0.99). (B) Scatter plot comparing the gene expression profiles between two biologically independent samples of *Dux*-activated control mESCs (Pearson correlation, r = 0.98). (C) Bar plot showing the $-\log_{10}$ [P value] of the gene ontology (GO) terms enriched in each category of genes (right, right-tailed Fisher's exact test). The number of genes in each category is indicated at the top of each GO enrichment plot. (D) Average occupancy plots and heatmaps of Zfp281 signal within 5 kb of the center of TSS (Transcription Start Sites) region of the 2C-regulated genes. (E) A box plot showing the $\log_2$ FC (fold change) of Zfp281-unbound-2C-upregulated transcripts in Zfp281-perturbed and control mESCs after *Dux* induction. The black central line is the median, the box limits indicate the upper and lower quartiles. (A-E) The X in sgX refers to the gene that sgRNA targets to; sgSv40 is negative control. P values were calculated by the Wilcoxon rank-sum test unless otherwise stated, ** < 0.01. The dots represent outliers.

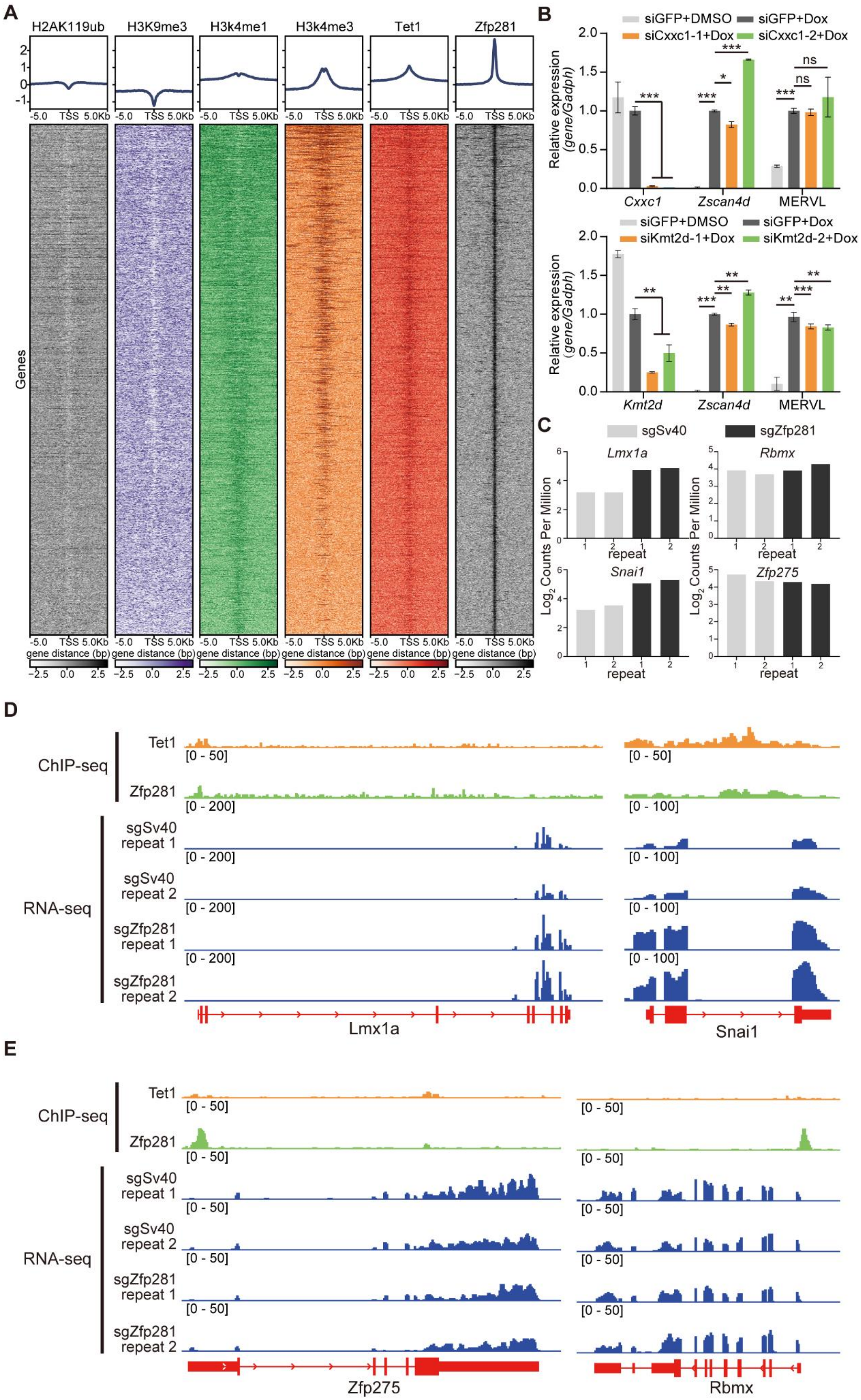

A
H2AK119ub
H3K9me3
H3k4me1
H3k4me3
Tet1
Zfp281
-5.0 TSS 5.0Kb
Genes
gene distance (bp)
B
siGFP+DMSO
siGFP+Dox
siCxxc1-1+Dox
siCxxc1-2+Dox
Relative expression (gene/Gadph)
Cxxc1
Zscan4d
MERVL
siKmt2d-1+Dox
siKmt2d-2+Dox
Kmt2d
C
sgSv40
sgZfp281
Lmx1a
Rbmx
Snai1
Zfp275
Log2 Counts Per Million
repeat
D
ChIP-seq
RNA-seq
sgSv40 repeat 1
sgSv40 repeat 2
sgZfp281 repeat 1
sgZfp281 repeat 2
Lmx1a
Snai1
E
Zfp275
Rbmx

**Fig. S3, Tet1 mediates the transcriptional regulation of Zfp281 on 2C-regulated genes.** (A) Average occupancy plots and heatmaps of the binding profiles in mESCs for Zfp281, Tet1, H2AK119ub, H3K9me3, H3K4me1, and H3K4me3 within 5 kb of the center of the Zfp281 peaks are shown. (B) Relative mRNA levels of *Cxxc1*, *Kmt2d*, *Zscan4d*, and MERVL normalized to *Gadph* in mESCs in mESCs upon indicated manipulation. Dox represents doxycycline. (C) Expression levels of *Lmx1a*, *Snail1* (Tet1 and Zfp281-cobound genes), *Rbmx*, and *Zfp275* (Zfp281-bound Tet1-unbound genes), in two biologically independent samples of Zfp281-perturbed or control mESCs after *Dux* induction. (D) RNA-seq and ChIP-seq genome browser track showing increased expression of the *Lmx1a* and *Snail1* (Zfp281 and Tet1-cobound genes) upon Zfp281 perturbation in Dux-activated mESCs. (E) RNA-seq and ChIP-seq genome browser track showing unchanged expression of the *Rbmx* and *Zfp275* (Zfp281-bound Tet1-unbound genes) upon Zfp281 perturbation in Dux-activated mESCs. The RNA-seq results are displayed as reads per kilobase million. (B-E) The X in siX/sgX refers to the gene that siRNA/sgRNA targets; siGFP and sgSv40 are the negative control. Shown are mean ± s.d, n = 3. P values were calculated by unpaired t-test, two-tailed, two-sample unequal variance, ns = no significance, * < 0.05, ** < 0.01, *** < 0.001.

#### Supplementary Table

**Table S1. Primers for real-time PCR**

| Gene | Sequence (5’-3’) |
|---|---|
| *Tet1* | F:- ACACAGTGGTGCTAATGCAG-<br>R:- AGCATGAACGGGAGAATCGG- |
| *Cxxc1* | F:- TTGGATGTGACAACTGCAACG-<br>R:- GTGGCGGTAACGAATCTCCAG- |
| *Kmt2d* | F:- GTGGCTGTTCCACACCCAG-<br>R:- AGCTTGAGCTTCTCAGCATCG- |
| MERVL | F:-CTCTACCACTTGGACCATATGAC-<br>R:-GAGGCTCCAAACAGCATCTCTA- |
| *Zfp352* | F:-AGAGGACAAGACCCAGTGCAG-<br>R:-GAGGTCCTCATCTGACCCAAG- |
| *Gapdh* | F:-CATGGCCTTCCGTGTTCCTA-<br>R:-GCCTGCTTACCACCTTCTT- |
| *Zscan4d* | F:-AAATGCCTTATGTCTGTTCCCTATG-<br>R:-TGTGGTAATTCCTCAGGTGACGAT- |
| syn*Dux* | F:-TCACAACCCCCACAAGTGG-<br>R:-TCCTCCAACTGTACTTCAGTAA- |
| *Dux* | F:-CAACGCAGGCTCTATGGAAC-<br>R:-GCTTCTTCCTGTGGCCAAAA- |

**Table S2. The sequence of the sgRNA**

| Gene | Sequence (5’-3’) |
|---|---|
| Zfp281-1 | GAGGATAACACGCACTGCGG |
| Zfp281-2 | GACTGATGAGCACGGCAACC |
